# Menstrual cycle phase modulates causal connectivity in the resting-state brain of healthy females

**DOI:** 10.1101/2024.06.07.598022

**Authors:** J. Mcleod, S. Sattari, A. Chavan, L. A.M. Galea, S. Babul, N. Virji-Babul

## Abstract

**Background:** Ovarian hormones exert direct and indirect influences on the brain; however, little is known about how these hormones impact causal brain connectivity. Studying the female brain at a single time point may be confounded by distinct hormone phases. Despite this, the menstrual cycle is often overlooked. The primary objective of this pilot study was to evaluate resting-state causal connectivity during the early follicular and mid-luteal menstrual phases corresponding to low *vs* high estradiol and progesterone, respectively.

**Methods:** Fourteen healthy control females (*M* = 20.36 years, *SD =*2.02) participated in this study. Participants were scheduled for two resting-state electroencephalography (EEG) scans during their monthly menstrual cycle. A saliva sample was also collected at each EEG session for hormone analyses. Causal connectivity was quantified using information flow rate of EEG source data. Demographic information, emotional empathy, and sleep quality were obtained from self-report questionnaires.

**Results:** Progesterone levels were significantly higher in the mid-luteal phase compared to the early follicular phase (*p* = .041). We observed distinct patterns of causal connectivity along the menstrual cycle. Connectivity in the early follicular phase was centralized and shifted posteriorly during the mid-luteal phase. During the early follicular phase, the primary regions driving activity were the right central and left/right parietal regions, with the left central region being the predominant receiver of activity. During the mid-luteal phase, connections were primarily transmitted from the right side and the main receiver region was the left occipital region. Network topology during the mid-luteal phase was found to be significantly more assortative compared to the early follicular phase.

**Conclusions:** The observed difference in causal connectivity demonstrates how network dynamics reorganize as a function of menstrual phase and level of progesterone. In the mid-luteal phase, there was a strong shift for information flow to be directed at visual spatial processing and visual attention areas, whereas in the follicular phase, there was strong information flow primarily within the sensory-motor regions. The mid-luteal phase was significantly more assortative, suggesting greater network efficiency and resilience. These results contribute to the emerging literature on brain-hormone interactions.

## Background

The brain’s interaction with sex steroid hormones, particularly through the menstrual cycle, represents a complex rhythmic relationship that is often oversimplified and underestimated in research. Studies commonly assess the human female brain at a single time point, failing to account for hormonal fluctuations inherent to the menstrual cycle (1–5). This approach limits our understanding of dynamic hormone-brain interactions.

The menstrual cycle, consisting of at least three phases (follicular, ovulatory, and luteal), is characterized by cyclical changes in ovarian hormones. In the early follicular phase, both estradiol (E2) and progesterone (P4) are at their lowest. During the luteal phase, P4 levels rise and reach their highest concentration during the mid-luteal phase, accompanied by a secondary peak in E2. This transition ushers in the onset of menstruation, thereby resetting the cycle.

E2 and P4 modulate brain activity by binding to their respective receptors. E2 receptors (ERs) and P4 receptors (PRs) are distributed throughout cortical and subcortical structures, with the most dense expression found in the prefrontal cortex, hippocampus, amygdala, and cerebellum (6–10). The distribution of ERs/PRs underscores the widespread influence that these hormones have on brain structure and function. Evidence from rodent (11–17) and human neuroimaging studies (18–21) have demonstrated dose- and time-dependent properties of E2 and P4 to modulate neurogenesis, myelination, and synaptic plasticity (for comprehensive review, see: (22–24)). The functions of E2 and P4 are made more complicated by their intricate interactions with each other. Initially, P4 synergizes with E2 to enhance its effects, but over time, P4 can actually diminish the effects of E2 (25–28). Adding additional complexity, E2 and P4 are only two hormones among many in the body’s endocrine system and do not act in isolation; their functions are influenced by the presence and levels of other hormones within the body (i.e., HPA axis).

Human female studies have demonstrated changes in both brain structure and function in relation to short-term hormone fluctuations (see (29) for a review). Specifically, positive associations between E2 and hippocampal gray matter volume, connectivity, and white matter integrity have been reported (6,30–34). Likewise, P4 is associated with increases in grey matter volume in the right basal ganglia (35), the hippocampus (36), and the subiculum (30).

Studies on human resting-state functional connectivity reveal that neuronal oscillatory patterns also shift in accordance with the menstrual cycle (see (37) for a review). Overall, E2 and P4 have been positively associated with functional connectivity across both hemispheres, within the cortex, and between cortical and subcortical regions, particularly within/between the DMN and SN (6,8,31,33,38–45).

The early follicular phase is associated with increased connectivity, between the midcingulate cortex and amygdala, the temporal and prefrontal cortex (45), and between the salience network (SN) and the posterior-default mode network (8). Recently, a single-subject, densely sampled study by Pritschet et al (38) demonstrated that whole-brain functional connectivity positively correlates with fluctuations in both E2 and P4 throughout the menstrual cycle, with the ovulatory surge in E2 having the most significant effect on connectivity. This peak in E2 has been repeatedly associated with an increase in connectivity within the default mode network (DMN) (8,33,39). During the mid-luteal phase, there is increased connectivity within the DMN, between the DMN and SN, from the anterior midcingulate cortex to the temporal-DMN, and between the hippocampus and the prefrontal and sensorimotor cortices. This phase also shows increased connectivity within the cingulate, insular, temporal, left angular gyrus, occipital, and cerebellar regions (8,46).

Although functional connectivity describes the correlation of activity between brain regions, it does not reveal how one region influences or causes activity in another. Directionality, as measured by causal connectivity, is an important characteristic of neural connections but has been less studied (47). To our knowledge, only two studies have examined causal connectivity throughout the human menstrual cycle – both of which have used fMRI (2,25).

Hidalgo-Lopez et al (48) evaluated causal connectivity in 58 naturally cycling healthy females and found that high-hormone phases exhibited greater connectivity within and between different brain networks. Specifically, during the early follicular phase, right-lateralized connections transmitted from the SN and DMN, with increased connectivity within the DMN, from the DMN to the executive control network (ECN), and from the parietal-ECN to the SN. The ovulatory surge in E2 was associated with greater connectivity from the left insula to the frontoparietal region and a decrease between the left and right parietal. In the mid-luteal phase, there were right-lateralized connections from the SN, and increased connectivity from the right insula to the frontal regions, from the frontal-ECN to the SN, and from the posterior-ECN to the posterior-DMN. More recent work by the same group (49) corroborated these findings, showing that compared to the follicular phase, the luteal phase had increased connectivity within the DMN and SN, from the DMN to the SN, and from the ECN to the DMN. There was also a decrease in connectivity from the parietal-ECN to the DMN and from the parietal-ECN to the SN. Additionally, a lateralized connectivity pattern increased from parietal-ECN to frontal-ECN in the right hemisphere and decreased from frontal-ECN to parietal-ECN in the left hemisphere.

In this study, we applied information flow rate to resting-state EEG in order to compare the rate and pattern of causal connectivity in healthy females during two phases of the menstrual cycle using Liang-Kleeman’s directional information flow rate (IFR). IFR was first conceptualized by Liang (50) using the principles of information entropy and dynamical systems theory and was applied to calculate causality within time series data (51). Unlike other models of causal connectivity (i.e., granger causality), IFR does not assume stationarity, making it particularly suitable for EEG analysis. The information flow rate measures the transfer of information between time series at different locations, identifying the brain sources that transmit and receive information. This data-driven quantitative method provides a network of brain connections, enabling the detection of causal links in the brain. Recently, IFR has been used to investigate connectivity associated with neuropathologies, such as concussion and autism spectrum disorder (52,53). This approach allowed us to investigate the influence of acute fluctuations in E2 and P4 on dynamic brain network communication.

## Methods

### Participants

Fourteen females participated in this study. The sample population comprised healthy individuals, predominantly varsity athletes, enrolled at the University of British Columbia. Inclusion criteria required all participants to be between 18 and 25 years, right-handed, not currently using hormone contraceptives, and have no history of neurological disorders. This study was approved by the University of British Columbia Behavioural Research Ethics Board-B (H22-00846). All participants gave informed written consent before enrolment.

### Study design

Cycle day at assessment was determined using each participant’s self-reported date of onset for their most recent period. Participants were scheduled for two EEG scans: once during their early follicular phase (cycle days 1-5) and again during their mid-luteal phase (cycle days 18-22). A saliva sample was collected at each EEG session for hormone analyses. The following questionnaires were also collected at each session: Sleep Condition Indicator (SCI), Pittsburgh Sleep Quality Index (PSQI), Toronto Empathy Questionnaire (TEQ), and a background questionnaire. The background questionnaire collected demographic data, including details about menstrual cycle patterns and consumption of alcohol, cannabis, nicotine, and medications in the preceding 12- and 24-hours.

### Saliva data collection and hormone assay

Estradiol and progesterone were evaluated in saliva samples collected using the Salivabio Passive Drool Collection Aid from Salimetrics. Participants were instructed to produce 1.0ml of unstimulated saliva. All participants self-reported adherence to the study protocol, which included abstaining from food, caffeine, sugar, and brushing their teeth for one hour before testing. After collection, samples were stored in a -80°c freezer until analysis.

Samples were assayed at the Salimetrics Salivalab (Carlsbad, CA) using Salivary Estradiol ELISA immunoassay (Cat. No. 1-3702) and Salivary Progesterone ELISA immunoassay (Cat. No. 1-1502). Each sample was measured in duplicate for each analyte. All inter- and intra-assay CV values ranged between 6.2% and 7.55%, indicating favourable test reliability.

### EEG data collection, preprocessing, and source reconstruction

Five minutes of resting-state eyes-closed EEG data was collected for each participant using a 64-channel hydrocel Geodesic Sensor Net connected to a Net Amps 400 high-impedance amplifier. Cz was the reference electrode during the recording setup. Data was collected at a 500 Hz sampling rate with scalp electrode impedances typically less than 50kΩ.

Raw EEG data was preprocessed using EEGLAB in MatLab (54), re-referenced to the average, and down-sampled to 250Hz. A Notch filter was applied to the data to remove the 60Hz line noise, followed by a bandpass filter between 0.5 and 50Hz. Non-brain artifacts identified through Independent Component Analysis and visual inspection were removed.

Clean EEG sensor data was reconstructed to source space using Brainstorm in MatLab (55). The ICBM152 brain template was used for the head model (56). The head model was divided into three sections (scalp, skull, and brain) for forward modeling. Forward modelling was constricted to the cortex. The inverse modelling method, Minimum Norm Estimate (MNE), and sLORETA were used to obtain the source space solution (57,58). The cortical surface was divided into ten regions of interest (ROIs) using the Desikan-Killiany atlas (see Figure 1D for details) (59).

**Figure 1.**
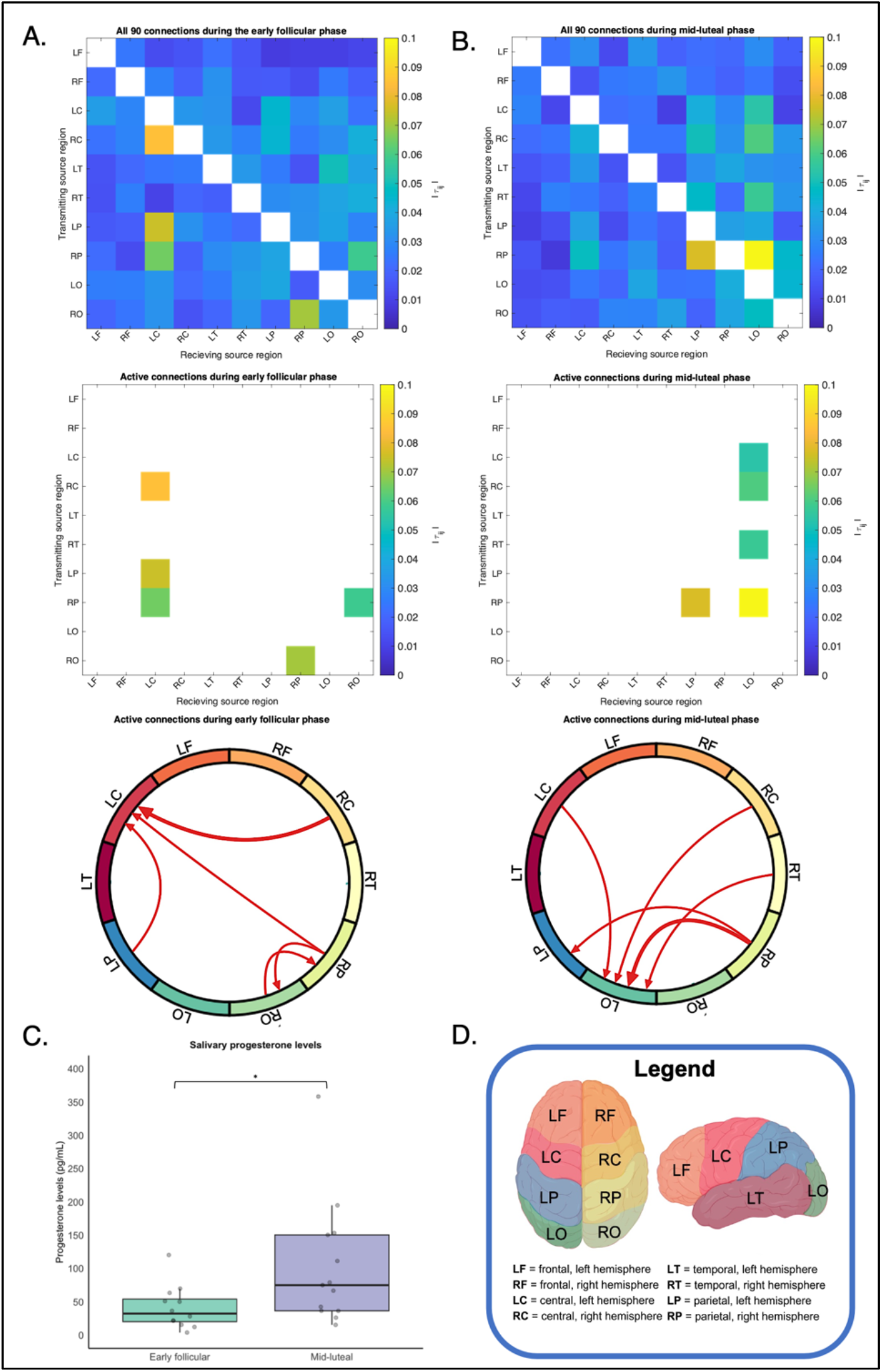
Causal connectivity during the (**A**) early follicular and (**B**) mid-luteal phases associated with (**C**) a statistically significant rise in P4 during the mid-luteal phase. Schematics A and B display, from top to bottom: binary matrix for all possible source pairs, binary matrix for active connections, and circular connectivity maps for active connections. Each matrix cell represents the mean absolute normalized information flow rates, |τi→j|, with source regions on the y-axis and receiving regions on the x-axis. The colour bar scale corresponds to |τi→j| values; darker colours indicate higher information transmission. Circular maps qualitatively show the spatial distribution of active connections. Arrows indicate active connections, with tails at transmitting regions and heads at receiving regions. The most active connection in each group is marked with a thick red arrow. (**D**) ROI source region legend colour-coded to the circular connectivity maps with definitions for abbreviated source regions used in A and B. * Denotes a statistically significant difference (p < 0.05).

**Figure 2.**
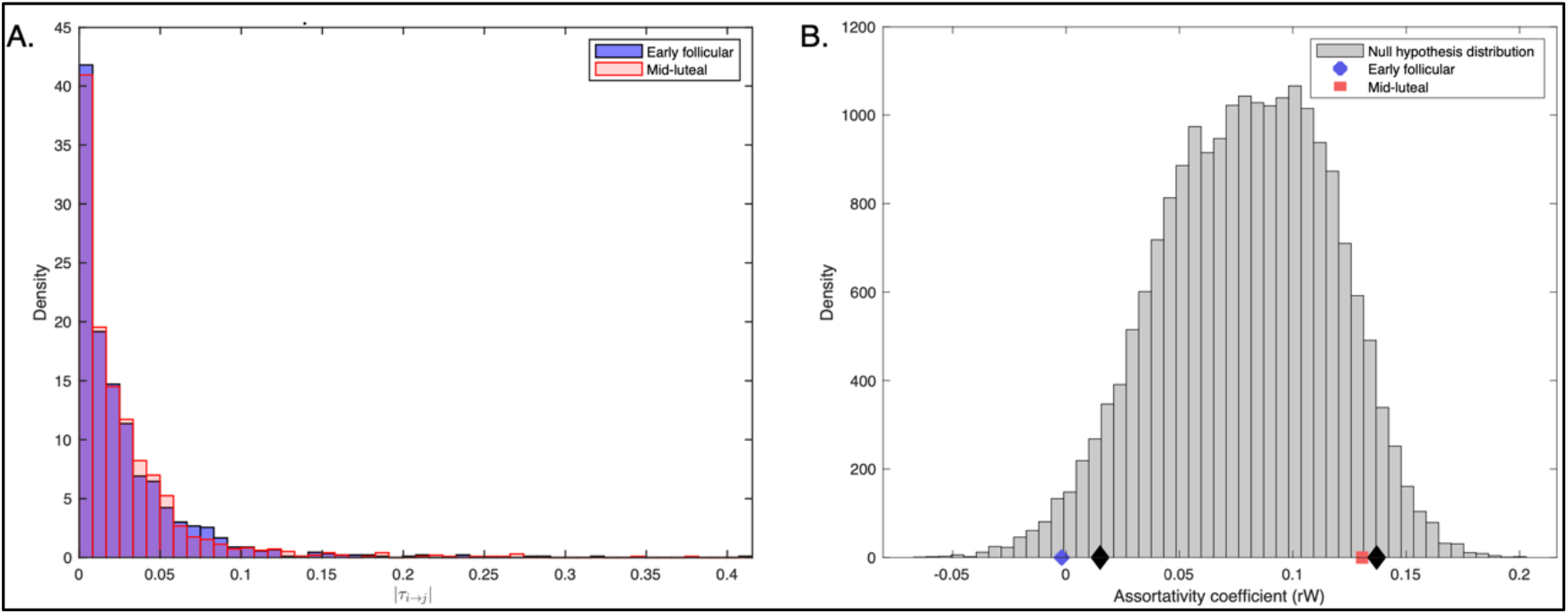
Distribution comparison of |τi→j| values and degree assortativity coefficients for the early follicular and mid-luteal phases. (**A**) Probability density histograms of the |τi→j| values for the early follicular (blue) and mid-luteal (red) phases, showing the overall magnitude of information flow across all possible connections. (**B**) Frequency histogram displaying the null hypothesis distribution of degree assortativity coefficient (rw) values (gray bars) alongside actual rw values for the early follicular phase (blue circle) and mid-luteal phase (red square). The black diamonds indicate the 5th and 95th percentiles.

### Causal connectivity

We calculated the normalized absolute information flow rate (|*τ*_*i→j*_|) between each possible pair of the ten source regions for each recording, resulting in 90 × 25 combinations. The information flow rate, *T*_*i*→*j*_, was derived using cross-correlation coefficients between the transmitter series, *v*_*j*_ , and the temporal derivative, *v′*_*j*_ of the receiver series, *v*_*j*_. This relationship is quantified by the *Liang-Kleeman coefficient* formula:

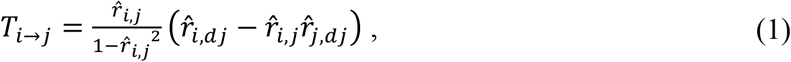

where 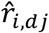 represents the correlation between *v*_*i*_ and *v′*_*j*_, and 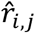 is the correlation between *v*_*i*_ and *v*_*j*_. The |*τ*_*i→j*_| is a value of coherence between ROIs; the larger the value of |*τ*_*i→j*_|, the more information that is transferred between the regions. This is also interpreted as an increase in the entropy of the receiver series as a result of information gained from the transmitter series. A connection between two source regions is considered *active* if the corresponding |*τ*_*i→j*_| exceeds the 0.05 threshold, suggesting that at least 5% of the receiver’s entropy rate is due to its interaction with the transmitter. See (60) for a comprehensive description of the information flow rate formula. All recordings during each phase were averaged to create a group mean, enabling the comparison of the magnitude of mean information flow rates and spatial patterns of the most active connections.

### Statistical analyses

Descriptive statistics (mean, standard deviation) were computed for the demographic data (ie., age, cycle length, cycle pattern). A linear mixed model was used to evaluate differences in levels of E2 and P4 across the two menstrual phases. The same model was also used to compare the Sleep Condition Indicator (SCI), Pittsburgh Sleep Quality Index (PSQI), Toronto Empathy Questionnaire (TEQ) scores, prior night’s sleep duration, and consumption within 12 and 24 hours prior to the study between the early follicular and mid-luteal phases. All tests were conducted at 5% significance. The analyses were completed using SPSS software (version 29.0.20.0).

The statistical significance of all connections was assessed using non-parametric permutation testing. For each individual in each group, the transmitter time series was permuted 100 times. This generated 100 permuted values for |*τ*_*i→j*_| for each connection. The significance threshold was set by calculating the 5th and 95th percentiles of the permuted distributions.

We performed a statistical analysis on all of the absolute normalized information flow rate values obtained from each group. Two distributions were generated from the |*τ*_*i→j*_| values that were calculated for each of the 90 pairs of 10 ROIs for each individual in both groups. The analysis was based on 12 × 90 = 1080 |*τ*_*i→j*_| values in the early follicular group and 13 × 90 = 1170 |*τ*_*i→j*_| values in the mid-luteal group. The Kolmogorov–Smirnov test was used to assess for differences in the shape of the distributions and the Kruskal–Wallis test was used to evaluate for differences in the central tendencies of the distributions.

The pattern of causal connectivity was statistically assessed using the degree assortativity coefficient (r_w_). The degree assortativity coefficient quantifies the Pearson correlation between the strengths of nodes at either end of an edge, indicating the tendency of nodes to connect with others of similar strength (61). It is a summary measure reflecting a network’s topological structure and is commonly used in brain network analyses. Mean r_w_ was calculated for each phase. Random permutations 10,000 times on the mean r_w_’s was performed to generate a frequency histogram from which the null distribution and actual r_w_ values were compared.

## Results

### Demographic and behavioural variables

Table 1 shows the descriptive statistics for all participants. Three subjects were lost to follow-up after participating in one session, resulting in *n*=11 participants undergoing repeated measures for each phase and *n*=3 undergoing single measures. In order to preserve power, the *n*=3 subjects undergoing single measures were not excluded from the analysis, resulting in *n=*12 measurements during the early follicular phase and *n=*13 measurements during the mid-luteal phase. The total participants sample had a mean age of 20 years and were predominantly of European ethnicity. Most participants reported monthly periods and a cycle length of approximately 30 days. There were no significant differences in self-reported behavioural measures (PSQI, SCI, TEQ) across the menstrual cycle.

**Table 1.**
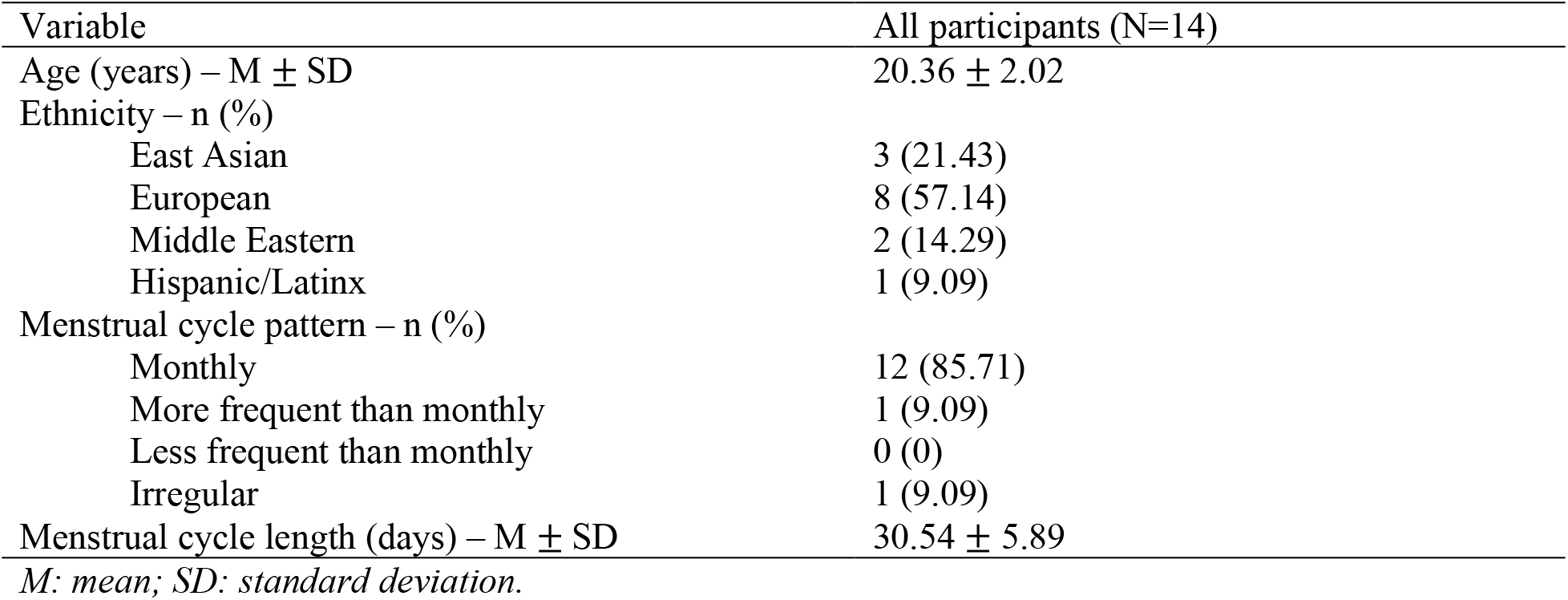
Descriptive statistics.

### Hormone levels

On average, assessment during the early follicular phase occurred on cycle day 3 and cycle day 20 for the mid-luteal phase. P4 levels averaged 41.14pg/ml in the early follicular phase and significantly increased to 103.47pg/ml in the mid-luteal phase (*p* = .041). Due to an error in the E2 assay, which led to invalid measurements, we removed it from the analysis. Results are shown in Table 2 and Figure 1C.

**Table 2.** Estradiol and progesterone levels measured during the early follicular and mid-luteal phases.

| | Early follicular ($n=12$) | Mid-luteal ($n=13$) | p-value |
| --- | --- | --- | --- |
| Cycle day of assessment | 3.83 ± 0.72 | 20.23 ± 1.48 | - |
| Progesterone (pg/ml) | 41.14 ± 32.46 | 103.47 ± 94.58 | .041* |

*Values are presented as mean ± standard deviation.*
*\* Denotes a statistically significant difference ( $p < 0.05$ ).*

### Causal connectivity

Information flow rate values were relatively equal between the two phases. The 90 mean |*τ*_*i→j*_| values ranged from 8.2567e-3 to 8.4992e-2 during the early follicular phase and from 7.8087e-3 to 9.8702e-2 during the mid-luteal phase. Five connections during each phase exceeded the *active* threshold (Table 3).

**Table 3.**
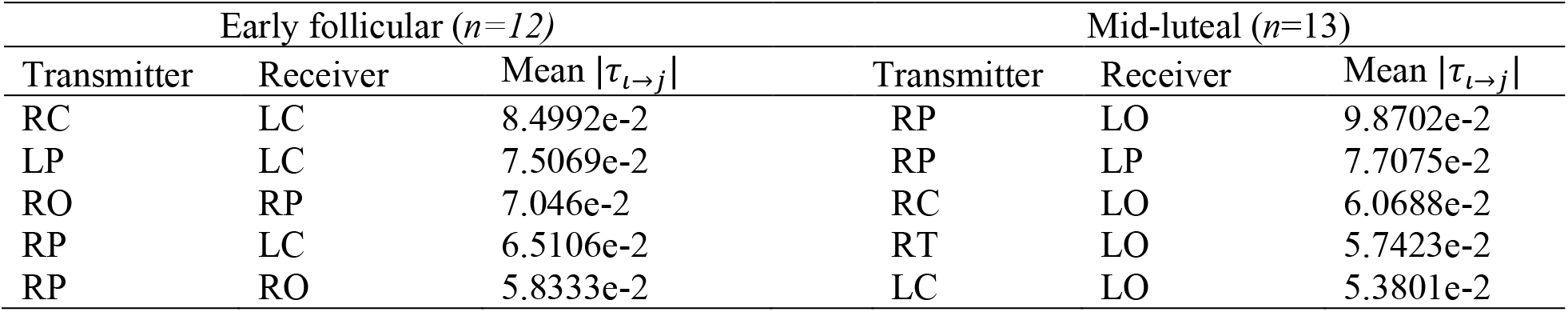
List of the active connections (|τ_*i*→*j*_| > 0.05) during early follicular and mid-luteal phase.

Information flow matrices are illustrated in Figure 1. We focused our qualitative comparison on the active connections during each phase. During the early follicular phase, activity was centralized. The primary regions driving activity were the right central and left/right parietal areas, with the left central region being the predominant receiver of activity. During the mid-luteal phase, activity was more posterior. Most active connections were transmitted from the right side, with the strongest two connections coming from the right parietal region and the left occipital being the main receiver.

### Statistical analysis of causal connectivity

All 90 mean connections in each phase were statistically significant. During the early follicular phase, the 5th and 95th percentiles were -1.551e-05 and 1.560e-05, respectively. In the luteal phase, these percentiles were -1.614e-05 and 1.624e-05. All mean connections exceeded the 95th percentile z-score, therefore, significance was confirmed.

The overall magnitude of information flow across all possible connections followed the same distribution during both phases (Figure 3A). This was confirmed statistically using the Kolmogorov–Smirnov and Kruskal-Wallis tests, which indicated no significant difference in the shape (*p* = 0.8892) or central tendency (*p* = 0.6673) between the two distributions, respectviely.

The pattern of connectivity significantly differed between the two phases. The mean degree assortativity coefficient was r_w_ = -0.00193 during the early follicular phase and r_w_ = 0.13058 during the mid-luteal phase. As illustrated in Figure 3B, the early follicular phase’s r_w_ was below the 5th percentile of the null hypothesis distribution. Therefore, we conclude that the mid-luteal phase is significantly more assortative than the early follicular phase.

## Discussion

In this preliminary study, we investigated resting-state causal connectivity across two time points in the menstrual cycle (early follicular and mid-luteal) in a sample of 14 healthy naturally cycling females. We applied information flow rate to resting-state EEG source data and generated two binary matrices of mean absolute normalized information flow rates between 90 unique pairs of ten ROIs. Overall, we observed distinct patterns of causal connectivity in the early follicular and mid-luteal cycle phases, which were associated with differences in salivary P4 levels.

As expected, P4 levels were significantly higher in the mid-luteal phase compared to the early follicular phase. Levels were within the same range reported by previous studies (43,48). This shift in P4 levels across the two phases was associated with distinct changes in causal connectivity. The early follicular phase was associated with a centralized connectivity pattern with the left frontal region as the primary receiver of activity, mainly recruited from the left/right parietal and right central/sensory-motor regions. In the mid-luteal phase, the connectivity pattern shifted posteriorly, with most connections being transmitted from the right side in an overall anterior-to-posterior direction. Two ERP studies found elevated P4 to be associated with increased activity in posterior brain regions (62,63). Similar to our findings, Haraguchi et al (63) found the first phase of the menstrual cycle to have greater activity in the frontal brain areas. Our findings are also consistent with work by Hidalgo-Lopez et al (48,49), who reported increased recruitment from the right side during the mid-luteal phase. However, Hidalgo-lopez et al (48) found a right-lateralized pattern during the early follicular phase, which we did not observe. Their use of seed-based ROI analyses, in contrast to our whole-cortex atlas approach, likely contributes to the differences we observed in connectivity patterns.

The shift in connectivity patterns that we observed may reflect a P4-dependent mechanism of plasticity. Peterson et al (46) suggested that differences in functional connectivity across the menstrual cycle were more closely associated with P4 rather than E2. However, these findings are primarily correlational, and without multi-modal structural and functional analyses, the findings are limited. The connectivity changes from the early follicular to mid-luteal phase might be explained by the combined effects of E2 and P4, with structural changes driven by fluctuations in the E2/P4 ratio potentially driving the shifts in causal connectivity. The absolute value of E2 and P4 might be less relevant than their relative changes over the cycle; however, our interpretation of this topic is limited by the error in the estradiol assay. It might also be fruitful for future studies to sample additional phases of the menstrual cycle to provide a more detailed time-based analysis of directionality with relative E2 and P4 levels. These interpretations remain speculative, and further studies are needed to investigate the combined versus separate effects of E2 and P4.

During the early follicular phase, three strong connections converge on the left central *(sensory-motor)* region from the right central *(sensory-motor)* and bilateral parietal regions. In contrast, during the mid-luteal phase, the most active connections are directed toward the left occipital *(visual)* region and originate from more widespread cortical areas: central, temporal, and parietal regions. Functionally, information flow during the early follicular phase is directed toward sensory-motor regions; however, in the mid-luteal phase, information flow shifts to areas responsible for visual-spatial processing. Various ERP studies have shown that elevated P4 during the mid-luteal phase is associated with improved visual memory, longer visual attention, quicker visual processing and perception, increased attentional blink, and greater visual-evoked responses (63–67). Findings from this study align with previous literature, indicating that the mid-luteal phase, and associated high levels of P4, is associated with increased activation in visual-perceptual processing areas. Interestingly, we observe these strong patterns in the resting-state while participants had their eyes closed. This highlights an important area for further investigation into the cyclical nature of visual processing. Also, the inclusion of a third measurement during the pre-ovulatory phase is likely to provide greater insights on the influence of the E2 surge on causal connectivity and help to differentiate the combined versus individual effects of E2 and P4.

We also observed a significant change in the topological structure of the network. Specifically, the degree assortativity coefficient was significantly higher during the mid-luteal phase compared to the early follicular phase. A high degree assortativity coefficient is indictive of a network characterized by clusters of highly connected nodes, where nodes preferentially connect with others of similar degree, forming dense sub-networks. In contrast, a low degree assortativity coefficient occurs when a network exhibits a disassortative pattern, with high-degree nodes more likely to connect with low-degree nodes, resulting in a more random and less clustered structure. A high degree assortativity coefficient is thought to reflect greater network efficiency and resilience (68). The neuroprotective properties of P4 offer a possible explanation for increased resilience. Interestingly, Andreano et al (8) suggested the mid-luteal phase to be a time of increased vulnerability, contrasting with our findings. They examined incidence rates for affective disorder across the menstrual cycle, reporting increased functional connectivity during the mid-luteal associated with heightened stress reactivity and memory for negative experiences. Importantly, they did not employ the degree assortativity coefficient, which could account for the differing conclusions. This discrepancy highlights the importance of considering different methodological definitions of resilience and vulnerability in the context of brain connectivity.

This study has several limitations. First, the error in the E2 assay interfered with our ability to draw conclusions about the role of E2 in causal connectivity. E2 and P4 function in a dynamic interplay, and the mid-luteal peaks in E2 and P4 likely represent a period of intricate causal connectivity changes resulting from their interaction. Future studies could incorporate more detailed analyses of different hormones as well as more timepoints to explore the richness of the different phases of the menstrual cycle.

Second, we only included females who were not using hormonal contraceptives, a common exclusion in menstrual cycle research. Hidalgo-Lopez (49) is, to our knowledge, the only group to consider menstrual cycle causal connectivity without excluding those on birth control. As of 2019, almost half of all women of reproductive age (15 to 49 years) worldwide (>300 million) were using hormonal contraceptives (69). Therefore, a significant portion of the mostly female population is overlooked in current research. In a follow-up study, we plan to recruit a sample of females using hormonal contraceptives, controlling for type and duration of use, to evaluate the influence of hormonal contraceptives on EEG causal connectivity. Additionally, we used a fixed generalized model of calendar counting to predict menstrual cycle phase, which overlooks individual variation in the menstrual cycle. Although out of scope for this study, longitudinal tracking of participants’ cycles prior to study enrollment could provide more personalized data that allows for more accurate phase measurements. Adding objective markers of menstrual phase (e.g., ovulation detection kits, ultrasound imaging of follicles) would also support more accurate phase measurements. Although salivary assays provided objective support for hormone levels at the time of EEG, using serum would enhance reliability, even though salivary levels report free hormone and serum measures both free and bound hormone levels. Another limitation is that we did not perform test-retest reliability for the causal connectivity analysis. Collecting a relatively short recording (5 minutes) limited our ability to complete a meaningful reliability test. In future studies, we intend to use longer recording durations.

## Conclusions

To our knowledge, this is the first study to use EEG to evaluate causal connectivity changes in healthy females across the menstrual cycle. The observed shifts in resting-state causal connectivity patterns highlight the potentially confounding effects of fluctuating progesterone and stress the need to consider cycle phase as a variables in research. Investigating these changes across the menstrual cycle is necessary for ensuring meaningful representation of females in research.

## Abbreviations

EEG: Electroencephalography
ROI: Region of interest
HRT: Hormone replacement therapy
ER: Estrogen receptor
PR: Progesterone receptor
DMN: Default mode network
SN: Salience network
ECN: Executive control network
IFR: Information flow rate
CV: Coefficient of variability
MNE: Minimum norm estimate
SCI: Sleep Condition Indicator
PSQI: Pittsburgh Sleep Quality Index
TEQ: Toronto Empathy Questionnaire
fMRI: Functional magnetic resonance imaging
DK: Desikan-Killiany atlas.

## Declarations

## Acknowledgements

We would like to thank all of the athletes from the University of British Columbia who participated in this study. Figure 1 was created with BioRender.com.

## Authors’Contributions

JM contributed to the study design, data collection, preprocessing, and analysis, as well as the writing of the manuscript. SS contributed to the data analysis. AC was involved in the development of the connectivity and source analysis pipeline. LG designed the study, contributed to the endocrine insights, and was involved in the writing of the manuscript. SB designed the study and was involved in the writing of the manuscript. NV-B designed the study, supervised the collection and processing of the data, developed the neurological insights, and contributed to the writing of the manuscript.

## Availability of data and materials

The datasets generated for this study are available on request to the corresponding author.

## Ethics approval and consent to participate

This study was reviewed and approved by the University of British Columbia Behavioural Research Ethics Board-B (H22-00846). All participants gave informed written consent before enrolment.

## Consent for publication

Not applicable.

